# Rapid Micronucleus-Sequencing (RaMN-Seq) Reveals Patterns of Bias in Induced and Spontaneous Chromosomal Instability

**DOI:** 10.64898/2026.10.14.759061

**Authors:** Jovanna K. Maharaj, Isabel McNeil, Molly A. Guscott, Thorsten Singer, Siegfried Hauch, Nischalan Pillay, Nadeem Shaikh, Sarah E. McClelland

## Abstract

Micronuclei are extra-nuclear structures formed following aberrant chromosome mis-segregation. Due to abnormal membrane features, micronuclei are liable to membrane rupture and are proposed as an important intermediate of extreme genome evolution events in cancer such as chromothripsis. Studying the features and contents of micronuclei can be informative about the underlying defective genomic instability process, but current purification strategies are time-consuming and require specialised methods. We developed <u>Ra</u>pid <u>M</u>icro<u>N</u>ucleus sequencing (‘RaMN-seq’) – a rapid (∼2 hour), kit-style micronucleus purification strategy coupled to single micronucleus low-pass DNA sequencing to explore the genomic contents of micronuclei. Using RaMN-Seq we recapitulate the known chromosome bias of nocodazole washout-induced chromosome mis-segregation in diploid cells. We explore the features of ruptured (cGAS-positive) and intact micronuclei, finding no substantial differences in chromosomal contents or sizes between rupture states. Finally, we extend RaMN-Seq to cancer models, demonstrating the ability to rapidly explore features of chromosomal instability in cancer cell lines and fresh-frozen tumour samples. Overall, RaMN-Seq allows simple, rapid purification of micronuclei from a range of sources, and offers an inexpensive alternative to single cell genome sequencing to examine properties and chromosomal identities of genetic material affected by chromosomal instability.

## Introduction

Genetic instability is a hallmark of cancer, manifesting on a scale ranging from single nucleotide changes to alterations in whole chromosome number and structure. Instability at the chromosomal scale – termed chromosomal instability (CIN) – includes events such as translocations of chromosome arms, aneuploidy due to aberrant chromosome number, and whole genome doubling (Lengauer, Kinzler and Vogelstein, 1998; Carter et al., 2012). CIN stems from both pre-mitotic and mitotic defects during cell division. Replication stress and unrepaired DNA damage lead to the formation of anaphase chromosome bridges and lagging, acentric chromosome fragments (Shaikh and Mazzagatti, 2022), while chromosome segregation defects during mitosis preferentially induce whole, lagging chromosomes (Thompson and Compton, 2008). Each of these processes can generate micronuclei (MN), membrane-encapsulated DNA-containing bodies separate from the interphase nucleus. Thus, the chromosomal content of MN can be indicative of the ongoing processes driving CIN in cells. Distinct drivers of CIN also impact the identity of chromosomes or fragments subject to chromosome mis-segregation and incorporation into MN, with previous studies demonstrating specific biases to mis-segregation (Worrall et al., 2018; Klaasen et al., 2022).

While micronuclei form spontaneously in cancer cell lines due to inherently high rates of CIN, these native micronuclei have rarely been analysed. Instead, micronuclei are frequently induced using aneugens such as nocodazole or reversine, which interfere with the mitotic apparatus and lead to whole chromosome mis-segregation, and clastogens such as hydroxyurea or aphidicolin, which increase the presence of chromosomal fragments via induction of DNA damage or replication stress (Hintzsche et al., 2017; Krupina, Goginashvili and Cleveland, 2021). These induction methods recapitulate the processes thought to drive CIN in cancer cells and, importantly, can be used on chromosomally stable cell lines to produce micronuclei, while avoiding the additional complexity of highly aneuploid cancer genomes. Nonetheless, the nature of micronuclear contents induced by such methods in diploid cells remains poorly studied, as the majority of studies have focused on cells with existing chromosomal instability (such as U2OS cells) treated with either nocodazole or reversine (Yasui et al., 2010; Crasta et al., 2012; Hatch et al., 2013).

Direct examination of micronuclear contents is not without its challenges. Due to their defective envelopes, micronuclei are more fragile, more prone to rupture, and less capable of repairing rupture than the primary nucleus. This, along with defective replication of DNA in micronuclei, contributes to their high levels of DNA damage (Crasta et al., 2012; Okamoto, Utani and Shimizu, 2012). Most damage present in micronuclei, as measured by γH2AX staining, occurs in S-phase or G2, with entry into S-phase seemingly required (Crasta et al., 2012; Hatch et al., 2013; Zhang et al., 2015). Micronuclear rupture can be measured by several markers, including the status of nuclear membrane proteins such as Lamin B1 and Lamin A/C, the loss of histone marks such as H3K27Ac, and the strong recruitment of cGAS (cyclic GMP-AMP synthase), which senses double-stranded DNA in the cytosol, to micronuclei once compartmentalization is lost (Hatch et al., 2013; Mammel et al., 2021; Mohr et al., 2021).

Currently, the primary method of micronucleus purification is based on sucrose gradient ultracentrifugation. Cells are lysed to separate the nuclear and cytoplasmic compartments, yielding a suspension composed of intact primary nuclei and micronuclei. Ultracentrifugation of this suspension results in separation of the micronuclei and nuclei by their relative densities. Thus, a relatively pure fraction of micronuclei is obtained, estimated to be approximately 80% of micronuclei from the cell culture, with 0.7% primary nuclei present (Abend et al., 1996). This approach can be combined with flow cytometry to specifically isolate ruptured micronuclei, for example, using fluorescently tagged cGAS to distinguish between ruptured and intact micronuclei (Toufektchan and Maciejowski, 2021). Recently, sucrose gradients have been employed without ultracentrifugation to purify micronuclei from peripheral blood lymphocytes (Wu et al., 2025).

Even without physical isolation, micronuclear genomic contents can be inferred through bioinformatic analysis of daughter cell pairs sequenced after micronucleus formation (Zhang et al., 2015). Nonetheless, purification and analysis of micronuclei remains key to gaining insights into their genomic, transcriptomic, and proteomic landscape. Some studies have directly sequenced micronuclei; Mic-Seq, utilized anucleated blood cells from MCM2-deficient mice, whose only genomic content are micronuclei formed in previous cell divisions, and identified genomic sites specifically and disproportionately affected by hydroxyurea treatment (Pruitt et al., 2017). Furthermore, both bulk and single micronucleus sequencing (MN-Seq) have been used to demonstrate the non-random patterns of aneuploidy in cancer cell lines, as well as a diploid cell line after induction of CIN (Klaasen et al., 2022). Additionally, a recent method exploits the use of laser microdissection to isolate micronuclei, which is highly precise but by necessity low-throughput and requires specialized equipment (Brückner et al., 2026).

Micronuclei therefore may be used to examine non-random aneuploidy patterns without the need for sequencing an entire cell. Thus, their contents may in turn provide an insight into the specific, current mechanisms of CIN at the time of micronucleus formation. Here we describe a simple and rapid method of purifying micronuclei, coupled with direct micronucleus (MN) sequencing (‘Rapid MN-Seq – RaMN-Seq’) in both bulk and single micronucleus approaches, applied to induced micronuclei from diploid cells as well as spontaneously forming micronuclei in chromosomally unstable cancer cell lines. We have also applied RaMN-Seq to purify and directly sequence micronuclei from fresh-frozen sarcoma tumour samples, underpinning the biological relevance and translational potential of the method. We demonstrate both the efficacy of our method as well as the utility of micronuclei as markers for ongoing chromosomal instability in both simple and complex genomic backgrounds.

## Results

### A filtration and benchtop centrifugation-based method for rapid and sensitive purification of micronuclei

We set out to develop a simple and rapid micronucleus purification method that circumvents the use of ultracentrifugation and instead utilizes a benchtop centrifuge and sequential filtration to separate micronuclei from nuclei based on size. Briefly, after harvesting, cells are resuspended in media containing cytochalasin B, which facilitates detachment of micronuclei from the primary nucleus via disruption of the actin cytoskeleton, before a brief set of lysis, centrifugation, and filtration steps are utilized to size-separate micronuclei from primary nuclei (**Figure 1a**). Importantly, to prevent nuclear deformation enabling passage of nuclei through the smallest filter size we added a light fixation step (see methods). In total, the method takes approximately two hours to obtain the final eluate containing micronuclei.

**Figure 1:**
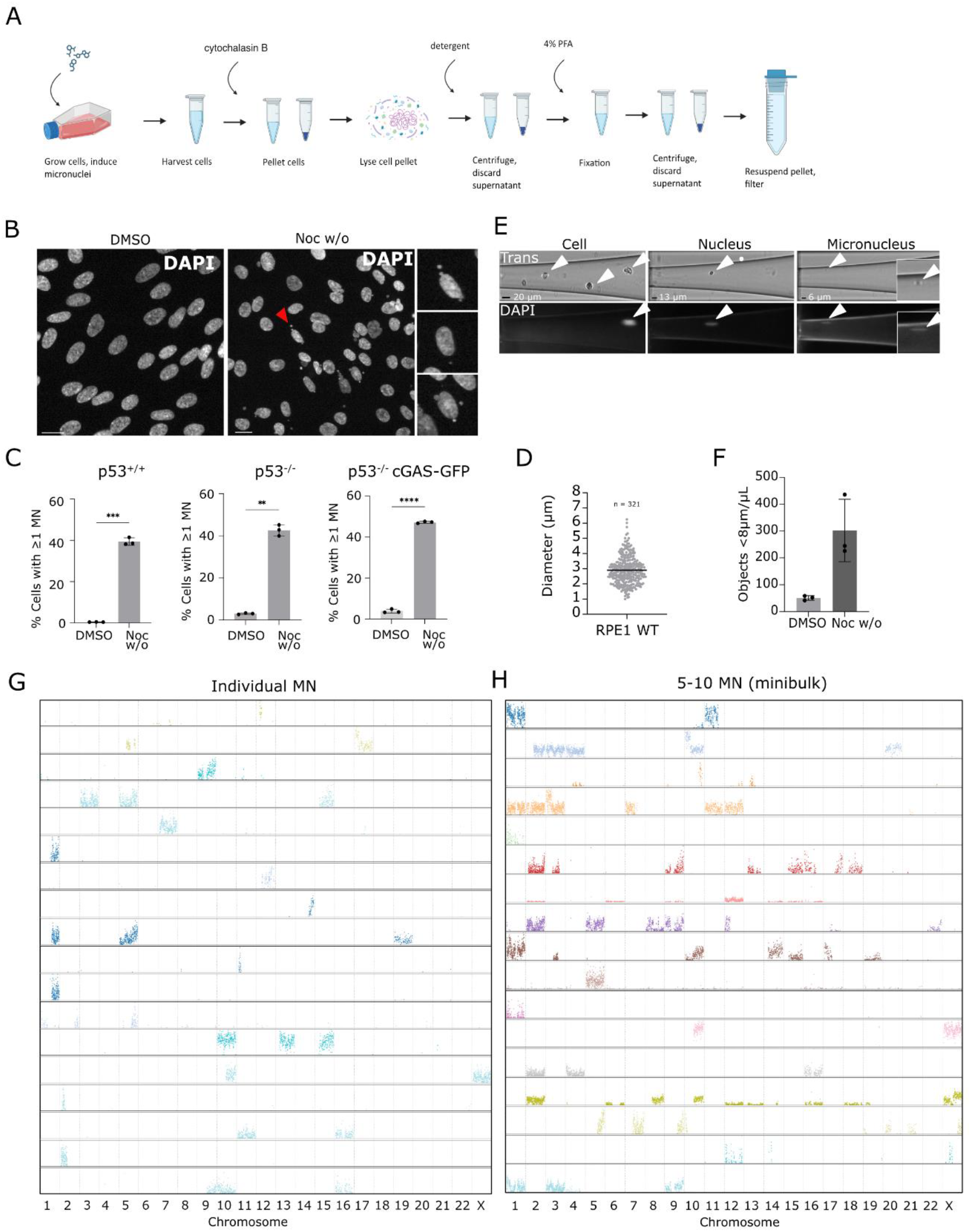
RaMN-Seq permits rapid, efficient isolation and sequencing of individual micronuclei induced by nocodazole washout. **A)** Schematic of RaMN-Seq purification protocol. Made with BioRender. **B)** Representative immunofluorescence images of RPE1 hTERT cells following nocodazole washout treatment. Scale bar = 20 µm. **C)** Percentage of cells with one or more micronuclei after nocodazole treatment. Mean and standard deviation of n = 3 biological replicates, 500 – 900 cells per replicate. **D)** Diameter of MN measured from immunofluorescence images, mean indicated, n = 3 biological replicates with > 100 MN scored per replicate. **E)** Representative images from CellenONE microfluidics nozzle indicating cell, nuclei, and MN in trans light and with 358 nm illumination fluorescence (DAPI). DAPI-positivity of cell (lower left panel) indicates a dead cell, whereas DAPI marks all nuclei and micronuclei since the cell membrane has been removed. **F)** Relative enrichment of particle sizes less than 8 µm with and without nocodazole treatment. DMSO n = 3 experiments; Noc w/o n = 3 (WT) experiments. **G)** Representative selection of single micronuclei sequenced after purification. Each row represents one single MN, with y-axis showing the read count for that segment of the genome. **H)** Representative selection of minibulked MN sequenced after purification, with each row representing a well with 5-10 putative MN.

To obtain cells with high rates of micronuclei we first treated non-transformed, hTERT-immortalized RPE1 cells and their p53^-/-^ cGAS-GFP counterpart with a nocodazole washout protocol which increases chromosome mis-segregation and micronucleus formation (Worrall et al., 2018). Note that the cGAS-GFP line was used for consistency with downstream experiments (see Figure 3), and the rate of micronucleus formation was similar between both RPE1 p53^+/+^ and RPE1 p53^-/-^ cGAS-GFP cell lines (**Figure 1b-d**). To characterise the micronuclei-containing eluate and evaluate the purity, we exploited the imaging capabilities of the CellenONE microfluidics device to measure the size of purified objects (**Figure 1e**). This revealed efficient enrichment for objects between smaller than 8 µm in nocodazole-treated cells that are rarely seen in control-treated RPE1 cells (**Figure 1f**) in line with their extremely low rate of spontaneous micronucleus formation (**Figure 1c**), suggesting these smaller objects are not debris or nuclear fragments resulting from the purification process. To confirm that these 4-8 µm-sized objects were micronuclei, we sorted individual objects using the CellenONE and performed low-pass genome sequencing using the DLP+ protocol (Laks et al., Cell 2019). Sequencing data revealed that the vast majority of objects < 8µm were individual micronuclei containing a low number of individual chromosomes/chromosome fragments (**Figure 1g**), with fewer than 1% nuclei observed, confirming the relative purity of our final eluate.

We note that the smallest object diameter accurately detected by the CellenONE platform is 3-4 µm, while immunofluorescence-based measurements of micronuclear size indicate a size range of 1 – 5 µm (**Figure 1d**), therefore we may not capture all micronuclei using CellenONE dispensation. We thus tested whether CellenONE-based dispensation could be bypassed using dilution seeding of the final filter eluate, which also permitted single micronucleus sequencing (**Figure S1**).

We wondered whether we could reduce library preparation and sequencing resources by combining multiple micronuclei in the same well before sequencing, in a ‘minibulk’ approach. To do this, we sorted 10 putative micronuclei per well from RPE1 WT cells after nocodazole washout and sequenced them in the same manner as with single RaMN-Seq. This approach reduced library preparation and sequencing costs ∼ten-fold and still enabled clear identification of micronuclear contents (**Figure 1h**).

### RaMN-Seq detects micronuclear chromosomal contents and recapitulates known biases in chromosome mis-segregation

Next, we verified that the purified and sequenced micronuclei recapitulated known features of nocodazole-induced micronuclei. We estimated the proportion of micronuclei containing one or more whole chromosomes by immunofluorescence imaging, staining with CREST antibodies, which recognise the centromere proteins CENP-A, B and C (Earnshaw and Rothfield, 1985). As expected, there was a strong bias towards whole-chromosome containing micronuclei after nocodazole treatment; only 5-10% of micronuclei were CREST-negative and therefore contained either a chromosome arm lacking the centromere, or a sub-chromosomal acentric fragment (**Figure 2a,b**). The presence of partial chromosomes encapsulated in micronuclei following induction of whole lagging chromosomes by nocodazole washout is likely due to an appreciable rate of damage to lagging chromosomes during cell division (Quignon et al., 2007). RaMN-seq largely reproduced this finding (**Figure 2c**), as the majority of micronuclei sequenced contained at least one whole chromosome.

**Figure 2:**
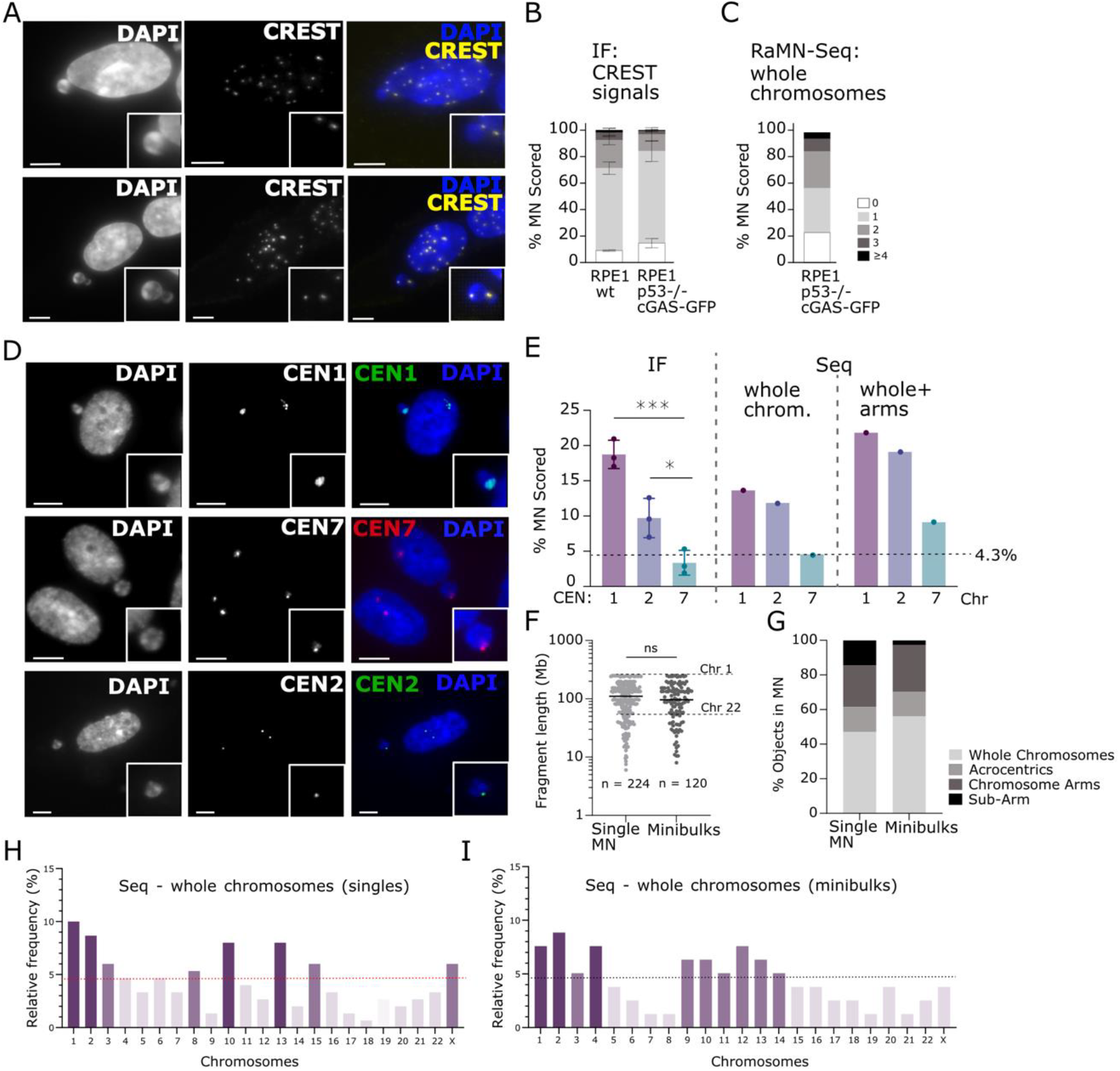
RaMN-Seq recapitulates chromosome mis-segregation biases. **A)** Representative immunofluorescence images of RPE1 cells 24h after nocodazole washout. DAPI (blue) indicates DNA, CREST (yellow) marks centromeres, scale bars = 5 μm. **B)** Quantification of centromere (CREST) foci per micronucleus in both RPE1 cell lines used, N = 3 biological replicates, >100 MN scored per experiment (WT total: 321; p53^-/-^ cGAS-GFP total: 339). **C)** Quantification of whole chromosomes (including acrocentric) per micronucleus from single MN sequencing in RPE1 p53 KO cGAS-GFP cells, n = 110 MN. **D)** Representative fluorescence *In-Situ* hybridisation (FISH) images of RPE1 WT cells probed for specific chromosome centromeres as indicated, CEN = centromere. Scale bars 5 um. **E) Left:** FISH quantification, n = 3 replicates, approx. 100 MN per replicate. **Right:** Quantification/MN frequency for whole chromosomes only based on sequencing (singles, RPE1 p53^-/-^ cGAS GFP). n = 110 MN for singles, or for whole chromosomes + chromosome arms only based on sequencing (singles, RPE1 p53 p53^-/-^ cGAS GFP). n = 110 MN for singles. **F)** Lengths in megabases of DNA (Mb) of individual fragments of DNA in MN. Mean object length indicated, Mann-Whitney test. N = 373 DNA fragments (from 110 single MN), N = 140 DNA fragments (from 24 minibulks). **G)** Quantification of chromosomal content of sequenced micronuclei from singles (n = 110) or minibulks (n = 24). Acrocentric chromosomes are counted separately. **H**,**I)** Histograms showing frequency of whole chromosomes in single MN (H) and minibulked MN (I).

Nocodazole washout induces preferential mis-segregation of chromosomes 1 and 2 due to a higher susceptibility to cohesion fatigue during a mitotic delay (Worrall et al., 2018). We assessed whether this bias is carried into micronuclear contents, first treating cells with nocodazole washout, and measuring the frequency of these chromosomes in micronuclei using fluorescence *In-Situ* hybridisation (FISH) against the centromeres of chromosomes 1, 2, and 7 (**Figure 2c,d**). In agreement with higher mis-segregation error rates (Worrall et al., 2018), chromosomes 1 and 2 were more frequently micronucleated than expected by chance (1/23 chromosomes = 4.3% by chance), while chromosome 7, here used as a control, was not (**Figure 2d, left**). Sequencing of single micronuclei confirmed this bias (**Figure 2d, middle**). The bias was even more pronounced when considering both whole chromosomes and chromosome arms (**Figure 2d, right**), suggesting the initial mis-segregation event of whole chromosomes can lead to micronucleation of chromosome arms and fragments. We saw no significant differences in terms of chromosome identity biases, sum genomic content, or numbers of chromosomes, arms or chromosome fragments within micronuclei whether single micronuclei or minibulks were analysed (**Figure 2e-h**). RaMN-Seq also revealed that chromosome 10 was micronucleated at a higher-than-expected rate (**Figure 2g,h**). Previous work (Worrall et al., 2018) also noted higher levels of chromosome 10 loss in RPE1 cells treated with nocodazole washout, supporting this observation. Overall, RaMN-Seq permits the selective purification of micronuclei that retain the features and chromosome identity observed using microscopy and/or FISH.

### Isolation of cGAS-positive and negative micronuclei reveals few differences between ruptured and intact micronuclei

Micronuclei accrue high levels of DNA damage and are considered a likely source of chromothripsis due to their deficiencies in replication and their propensity to rupture, exposing their DNA to the cytosol and leading to recruitment of cGAS (Hatch et al., 2013; Zhang et al. 2015; Mammel et al., 2021; Mohr et al., 2021). However, the precise mechanisms leading to micronuclear rupture, including whether certain features might predispose them to rupture, are not fully known. To investigate whether chromosome content identity could influence the likelihood of rupture, we used the RPE1-cGAS-GFP cell line (in the p53^-/-^ background) to distinguish between intact and ruptured micronuclei following nocodazole washout. We noted that at the timepoint where we purify micronuclei, ∼ 30% micronuclei were strongly cGAS-positive by immunofluorescence (**Figure 3a,b**). We also observed a significant amount of DNA damage, detected via γH2AX staining, with two distinct phenotypes present: Either a low number of distinct foci within the micronucleus, or intense, pan-MN staining (**Figure 3a, lower panel**). The levels and classes of DNA damage present were similar between ruptured and non-ruptured micronuclei (**Figure 3c**). Interestingly, when measured using immunofluorescence, cGAS-positive micronuclei had slightly but significantly smaller diameters than their cGAS-negative counterparts (**Figure 3d**).

**Figure 3:**
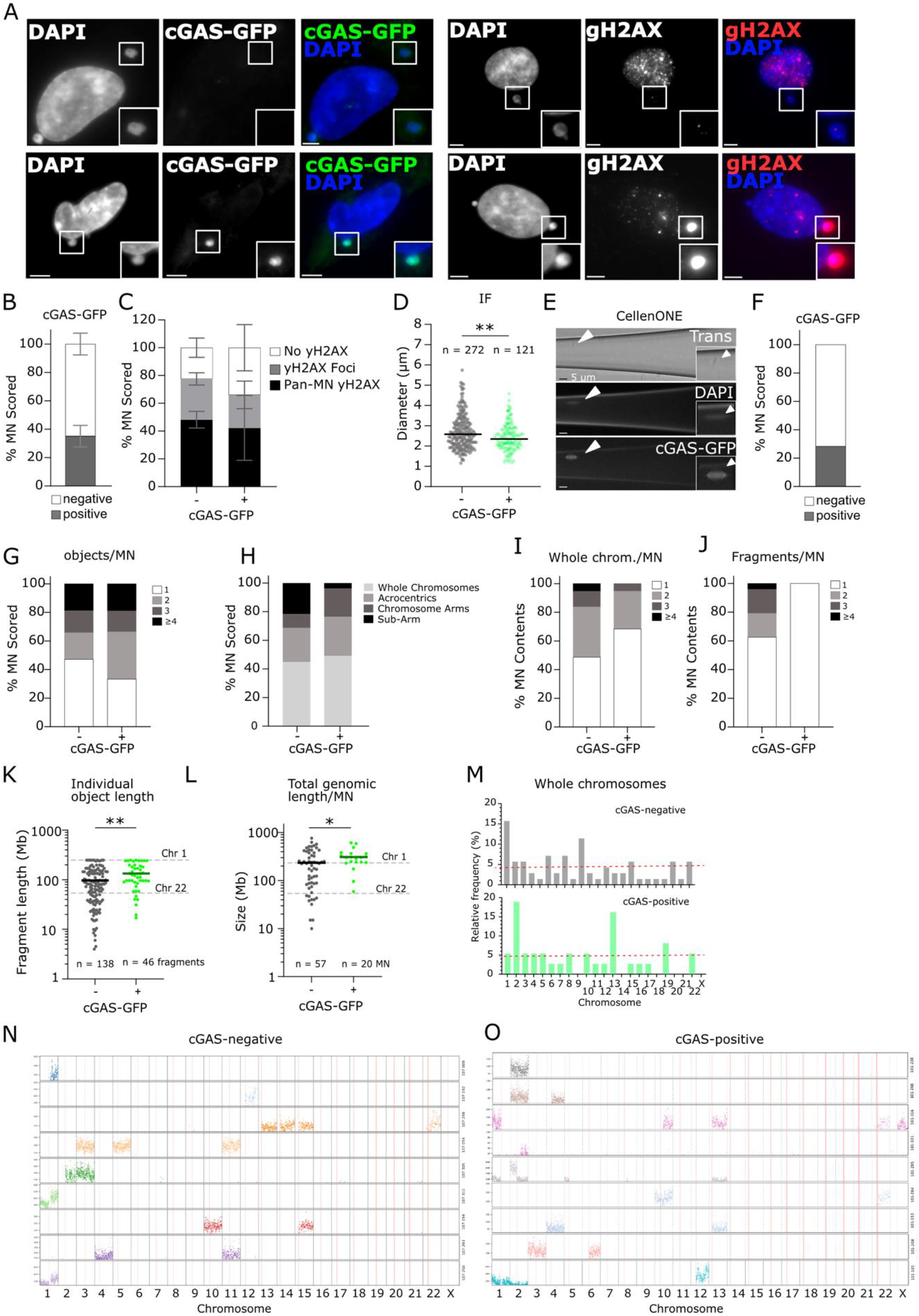
Analysis of ruptured and intact micronuclei. **A)** Representative immunofluorescence images of RPE1 p53^-/-^ cGAS-GFP, cells 24 h after nocodazole washout. DAPI (blue) indicates DNA, CREST (yellow) marks centromeres, red marks anti-*γ*H2AX antibodies. Scale bars = 5 µm. **B)** Proportion of cGAS-positive micronuclei after nocodazole treatment from immunofluorescence. **C)** Proportion of micronuclei (stratified into cGAS negative and positive) with γH2AX signal separated into foci or pan-micronuclear. **D)** Diameter of ruptured and intact MN after nocodazole washout measured from fixed cell immunofluorescence. Data in B and D from 3 biological repeats, approx. 100 micronuclei imaged per repeat. **E)** Images of CellenONE-based detection of cGAS-GFP-positive micronuclei. **F)** Proportion of cGAS-positive micronuclei identified with CellenONE imaging and sorting. **G)** Number of chromosome objects per micronucleus detected with sequencing, 81 micronuclei analysed. **H)** Chromosomal content of micronuclei detected with sequencing (acrocentrics counted as arms), from 1 repeat, 83 micronuclei analysed. **I**,**J)** Numbers of whole chromosomes (I) or chromosome fragments (J) in micronuclei detected with sequencing. **K)** Sizes of individual fragments of DNA in micronuclei. Mean shown, Mann-Whitney test, p = 0.002. N = 241 objects (cGAS-negative), 59 objects (cGAS-positive). **L)** Total genomic length in megabases of content of each micronucleus. N = 62 MN (cGAS-negative), n = 19 (cGAS-positive). Mean shown, Mann-Whitney test, p = 0.037. **M)** Percentage of each whole chromosome identified in micronuclei. N = 62 MN (cGAS-negative), 19 MN (cGAS-positive). **N**,**O)** Representative selection of single micronuclei sequenced after purification for cGAS-negative (N) and positive (O) micronuclei.

We performed RaMN-Seq on these cells after micronuclei were induced via nocodazole washout, measuring GFP fluorescence on the CellenONE to identify ruptured MN (**Figure 3e,f**). The proportion of ruptured micronuclei measured using the CellenONE was concordant with our immunofluorescence data (**Figure 3f**), and we were able to confidently separate micronuclei by their rupture status to investigate whether there were differences in chromosomal content between ruptured and intact micronuclei using low-pass sequencing. Ruptured micronuclei were slightly more likely to contain whole or multiple chromosomes and acrocentric chromosomes than intact micronuclei, and less likely to contain chromosome fragments (**Figure 3g-j**). There was also a small but significant difference between mean DNA fragment length in ruptured and intact micronuclei, with ruptured micronuclei more likely to contain larger chromosomal fragments (**Figure 3k**), and a slightly higher total genomic content (**Figure 3l**). In terms of chromosomal identities, intriguingly, chromosome 1 appeared enriched only in cGAS-negative (non-ruptured) micronuclei, while chromosome 2 was only enriched in cGAS-positive (ruptured) micronuclei (**Figure 3m-o**). A larger dataset will be required to determine whether this is a statistically significant observation.

### Aphidicolin treatment reveals distinct bias towards smaller chromosomal events

Treatment with aphidicolin is known to induce replication stress, leading to double-stranded DNA breaks and the mis-segregation and subsequent micronucleation of acentric chromosome fragments (Naim and Rosselli, 2009; Shaikh et al, 2022; Wilhelm et al, 2019). RPE1 p53^-/-^ cGAS-GFP cells were treated with 0.6 μM aphidicolin for 24 hours to induce micronuclei prior to harvesting and RaMN-Seq of single micronuclei. Micronuclei were induced in ∼ 10% cells (**Figure 4a,b**). In line with the expected increase of MN containing acentric chromosome fragments, RaMN-seq revealed a skew towards chromosome fragments (**Figure 4c**) compared to nocodazole washout (**Figure 2g**). This was also reflected in the sizes of individual DNA fragments present which were on average smaller than the smallest autosome (**Figure 4d**). Aphidicolin-induced micronuclei also contained overall less DNA, with an average of 100 Mb per micronucleus (**Figure 4e**), compared to 250-300 Mb in nocodazole-induced MN (**see Figure 2**).

**Figure 4:**
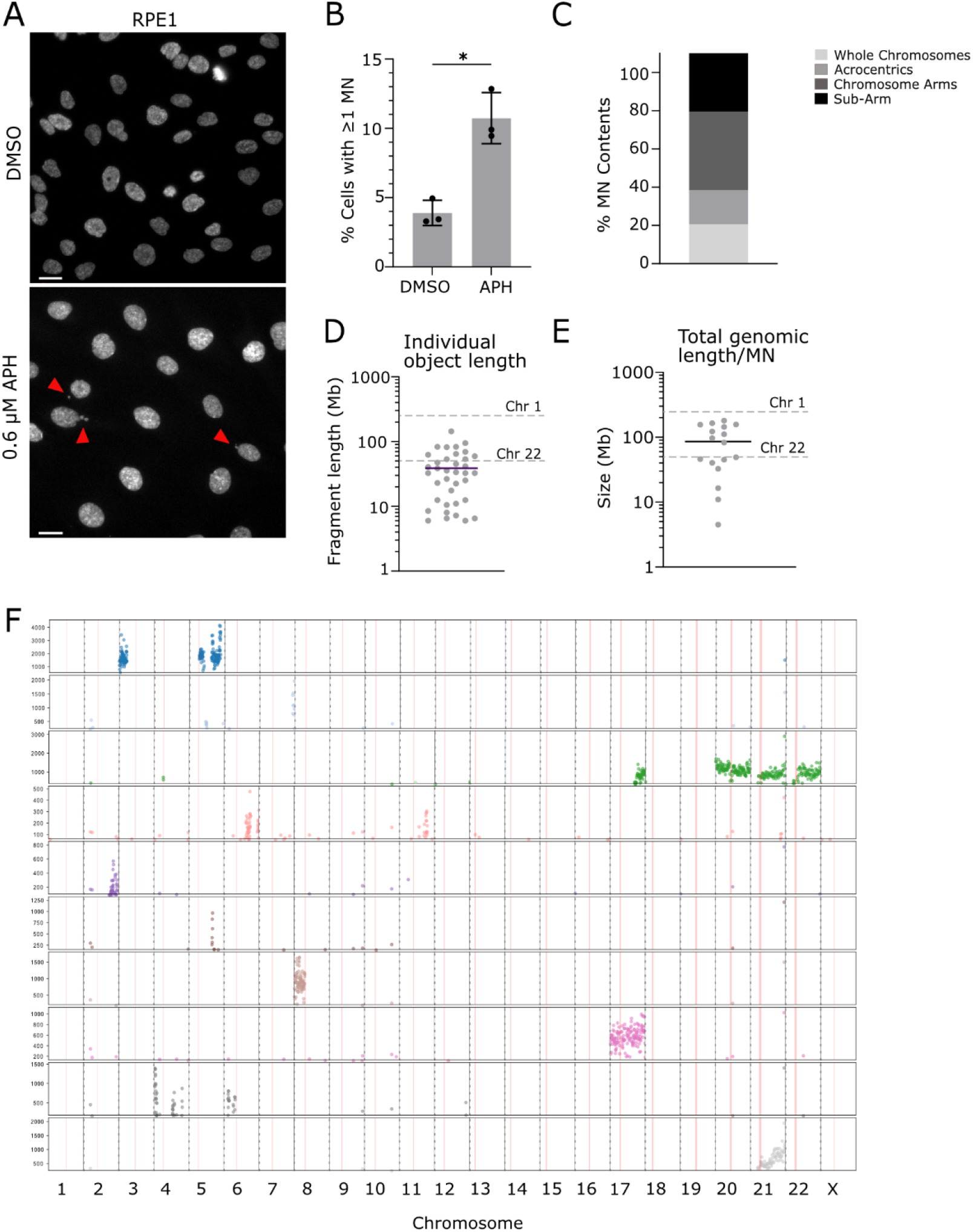
Analysis of aphidicolin-induced micronuclei. **A)** Representative immunofluorescence images of RPE1 p53^-/-^ cGAS-GFP cells after 24 h 0.6 μM aphidicolin. DAPI (grey) indicates nuclei, and red arrows indicate micronucleated cells. Scale bar = 20 μm. **B)** Micronucleus frequency after aphidicolin treatment. N = 3, 400-600 cells scored per biological repeat. Welch’s T-test, p = 0.011. **C)** Chromosomal content of micronuclei detected after sequencing from 2 repeats, n = 17 MN analysed. **D)** Sizes of individual fragments of DNA in micronuclei. N = 36 fragments from 17 MN. **E)** Total genomic length in megabases within each micronucleus. N = 17 MN. **F)** Representative selection of single micronuclei sequenced with RaMN-Seq.

### Application of RaMN-seq to cancer models reveals features of endogenous cancer CIN

Next, we applied RaMN-seq to cancer models. We first performed RaMN-Seq on a soft tissue sarcoma cell line LMS01 that exhibits a high level of endogenous chromosomal instability (∼ 15% cells with MN; **Figure 5a,b**). Using minibulked RaMN-seq, we observed a higher rate of chromosome arms and chromosome fragments compared to RPE1 cells treated with nocodazole-washout (**Figure 5e**) and noted that chromosome fragments were of a smaller size on average (**Figure 5f**). In terms of whole chromosome identities, we observed a strong bias towards chromosome 17 (**Figure 5g,h**). Chromosome arm identities were more distributed, though tending towards larger chromosomes (**Figure 5f**).

**Figure 4:**
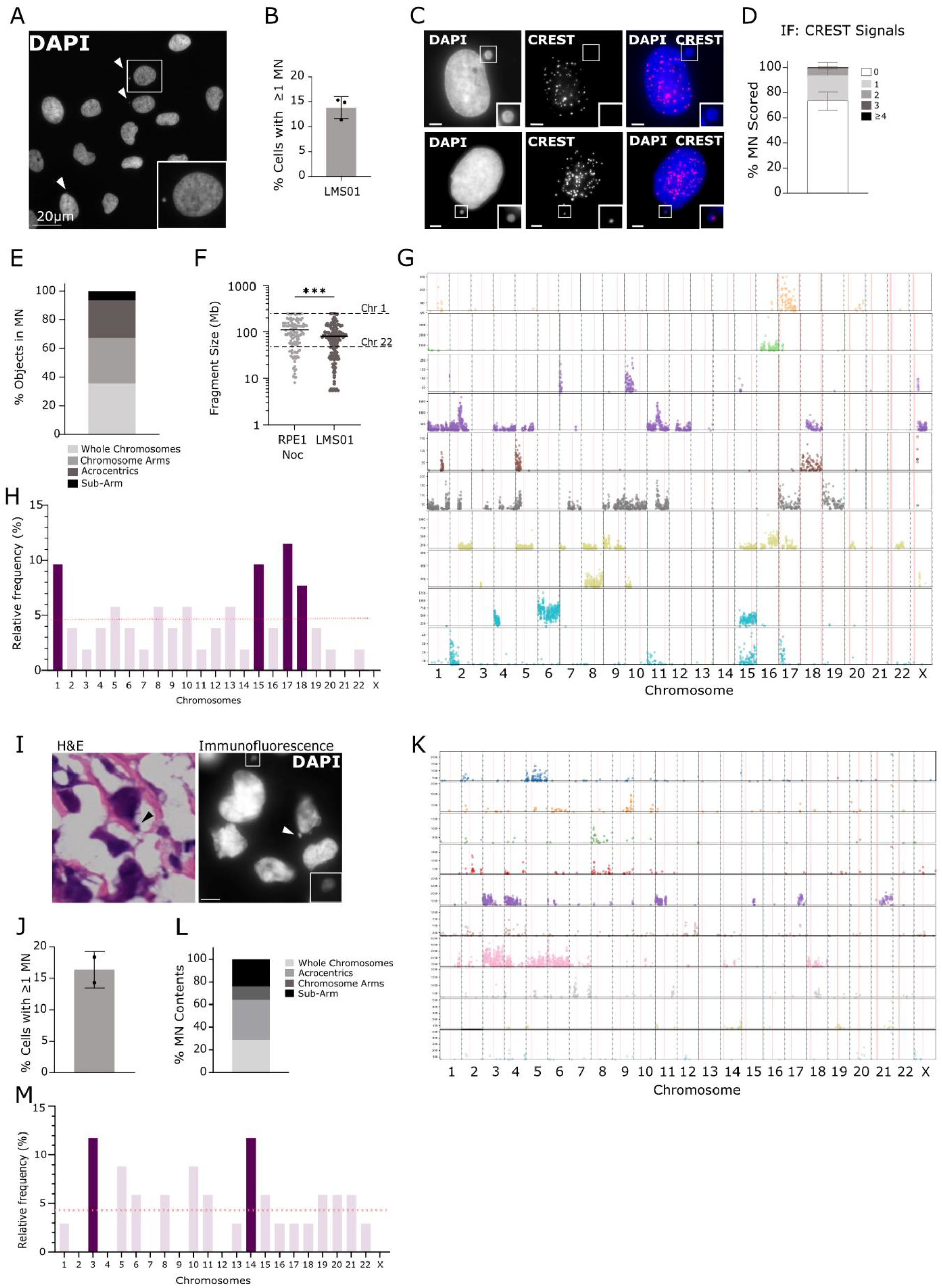
RaMN-seq is applicable to cancer cell lines and tumours. **A)** Representative immunofluorescence image of sarcoma cell line LMS01. **B)** Percentage of cells with micronuclei, n = 3 biological replicates, 500-900 cells scored per experiment. **C)** Representative immunofluorescence image of LMS01 with CREST (red) staining to indicate centromeres. **D)** Quantification of centromere (CREST) foci per micronucleus in LMS01 cells. N = 3 biological replicates, > 100 MN scored per replicate. **E)** Object types in MN. **F)** Fragment lengths within micronuclei, with nocodazole washout data from RPE1 cells reproduced from Figure 2 as a comparison. **G)** Representative RaMN-seq copy number profiles from micronuclei isolated from LMS01 cells (minibulks). **H)** Histogram of whole chromosome identities from RaMN-seq (minibulks). N = 27 minibulks (max 270 MN). **I)** Hematoxylin and Eosin staining (left panel) and immunofluorescence (right panel) images of sarcoma tumour sample 49538. DAPI (blue) indicates DNA. **J)** Percentage of tumour cells with micronuclei scored from immunofluorescence imaging. Two sections were scored, n= 628 and 679 cells from each. Both sections were then pooled for RaMN-Seq preparation. **K)** Representative RaMN-Seq profiles of putative MN isolated from tumour sample 49538. **L)** Chromosomal content of micronuclei detected with sequencing. **M)** Histogram of whole chromosome identities. N = 26 micronuclei.

We also tested whether RaMN-seq could be applied to tumour tissue. Sections of fresh-frozen sarcoma tumour samples were cut for both Haematoxylin and Eosin staining and immunofluorescence (**Figure 5i**), and parallel tissue samples were processed for RaMN-seq. Tumour 49538 was a histologically unclassified primary sarcoma, and matched bulk WGS sequencing data showed it was primarily diploid with low levels of genome alterations (data not shown; personal communication from N. Pillay). We quantified the rate of micronuclei using immunofluorescence, measuring a frequency of ∼16% (**Figure 5i,j**). We then purified and sequenced single micronuclei (**Figure 5k**). We noted a lower resolution due to lower sequencing depth and fewer micronuclei were sequenced than in previous samples, suggesting further optimization is required, and as a result we were unable to deconvolve MN contents below the level of chromosome arms. However, we observed overrepresentation of a distinct set of chromosomes compared to RPE1 and LMS01 cell line models, noting a particular bias towards chromosome 3 (**Figure 5j,k**).

## Discussion

We set out to design a simple, rapid, and inexpensive method to isolate and sequence the genomic contents of micronuclei. Using only benchtop equipment and commercially available reagents we can isolate micronuclei within two hours and identify their contents using single, or minibulk low pass genome sequencing. We verified the bias towards mis-segregation of chromosomes 1 and 2 in RPE1 cells treated with nocodazole washout and demonstrated that the contents of MN differ depending on the type of chromosomal instability induced. We explored whether micronuclear rupture was related to specific genomic contents. Overall, no major differences in content in terms of sizes or types of chromosome structures between ruptured or intact micronuclei were observed.

We also explored the utility of RaMN-Seq applied to cancer cell lines and a tumour fresh tissue sample. We reveal the feasibility of applying RaMN-Seq to fresh frozen tissue and demonstrated distinct sets of chromosome identity biases in cancer models compared to micronuclei induced with nocodazole washout. While this protocol is suitable for cancer cell lines with a wide range of micronuclei frequencies, RaMN-seq in tumour samples is likely limited to fresh or fresh-frozen tumours, due to the lower DNA quality in FFPE preparations. In this study we primarily used the CellenONE microfluidics device for dispensing micronuclei for downstream sequencing, but use of flow cytometry to perform the initial sorting step for single micronucleus sequencing, or serial dilutions, is also possible where a similar microfluidics device is not available. We primarily focused on sequencing single micronuclei, but use of minibulks consisting of up to 10 micronuclei still provided useful information regarding biases in micronuclear contents, particularly at a whole chromosome level, and may in fact be more accessible and more suitable for certain applications.

Overall, we have purified and sequenced micronuclei directly from a wide range of cellular and tumour models. With more data we will be able to deeply interrogate not just the chromosome identity of micronuclear contents, but frequent hotspots of breakage in the genome. Our work opens the possibility of utilizing micronuclei to scan for specific mechanisms of chromosomal instability in different cancer models and allows us to distinguish between ongoing and historical chromosomal instability.

## Methods

### Cell culture

The sarcoma cell line LMS01 was a gift from the University of Sheffield. LMS01 was cultured in Nunc™ EasYFlask™ cell culture flasks (ThermoFisher; #156499), with Gibco™ RPMI media (ThermoFisher; #11875093) supplemented with 10% (v/v) Gibco™ Foetal Bovine Serum (FBS) (ThermoFisher; #10500064), 1% penicillin-streptomycin (Sigma-Aldrich; #P4333), 1% (v/v) L-glutamine (Sigma-Aldrich; #59202c), 0.4 mg/mL L-glucose (Sigma-Aldrich; #G7021), and 5 μg/mL amphotericin B (ThermoFisher; #15290026). RPE1 p53 WT cells were from ATCC. RPE1 p53^-/-^ cells were a gift from the Godinho lab. RPE1 cells were grown in Gibco™ DMEM-F12 media (ThermoFisher; #11320033) with 10% FBS and 1% penicillin-streptomycin. HEK293T cells were grown in Gibco™ DMEM media (ThermoFisher; #11965092) with 10% FBS and 1% penicillin-streptomycin. All cells were maintained at 37°C and 5% CO_2_. Routine mycoplasma tests were conducted.

### Drug treatments

RPE1 cells were treated with 100 ng/mL nocodazole (Sigma-Aldrich; #M1404) for 16 hours, before a mitotic shake-off was performed, mitotic cells collected, pelleted, and then replated. Cells were allowed to recover for 24 hours prior to RaMN-seq, FISH, or immunofluorescence experiments. RPE1 cells were treated with 0.6 µM aphidicolin (Sigma-Aldrich; #A4487) for 24h prior to harvesting and RaMN-seq.

### Cell line generation

RPE1 p53^-/-^ cGAS-GFP cell lines were generated via lentiviral transduction of the pTRIP-CMV-GFP-FLAG-cGAS plasmid (Addgene #86675) into RPE1 p53^-/-^ cells. 10 μg of the cGAS plasmid was transfected into HEK293T cells alongside 3 μg pMD2.G (Addgene #12259) and 6.5 μg psPAX2 (Addgene #12260). DNA solution and transfection reagent (Lipofectamine 2000; ThermoFisher #11668027) were prepared separately in Opti-MEM™ media (Gibco; #31985062) and mixed. Transfection was performed by adding the mix dropwise onto the cell culture plates and left overnight. Media was changed to DMEM with 10% FBS without pen-strep to allow for viral production. Virus was harvested after 24h, mixed with 1/1000 polybrene (Sigma-Aldrich; #H9268-5G), and added to the recipient cells. Media on the recipient cells was changed after 24 hours, and cells were assessed for fluorescent signal after 48 hours. Recipient cells were then single-cell sorted and expanded to establish clonal populations. Clones were screened and one was selected for use in experiments based on a strong nuclear cGAS localisation and CIN rates similar to that of the parental cell line.

### Tumour sample processing

Fresh-frozen sarcoma tumour samples were cut into small pieces and treated with 100 ug/mL collagenase in Hank’s Balanced Salt Solution for 30 mins at 37°C, with shaking at 100 rpm for 20 minutes for enzymatic dissociation. They were homogenized to a single-cell suspension using a 100 µm cellStrainer filter and rinsed with FACS buffer (1% FBS and EGTA in PBS). This suspension was pelleted at 300 x g for 6 minutes, at 4°C, and the supernatant removed. Cells were resuspended in 2 mL RBC Lysis Buffer and incubated for 2 minutes at room temperature. FACS buffer was added to a volume of 50 mL, and the sample was centrifuged at 300 x g at 4° C for 6 minutes. The supernatant was aspirated, and the pellet was resuspended in FACS buffer, filtered once more, and centrifuged again under the same conditions. This wash step was repeated, and the resultant single cell suspension was used for the RaMN-Seq protocol.

### RaMN-Seq

Cells were harvested when 80-90% confluent or after any treatment, trypsinized, and collected in 1 mL media. Cytochalasin B (Sigma-Aldrich; #C2743) was added to the media for a final concentration of 10 µg/mL and cells were centrifuged at 21,000 g for 30 mins at 37°C. The supernatant was discarded, and the pellet was resuspended in 500 µL Lysis Buffer NL (from the Qiagen Qproteome Nuclear Protein Kit, Qiagen; #37582) supplemented with 0.1 M DTT and 5 µL protease inhibitors. Cells were lysed on ice for 15 mins before being transferred to a clean, cold Eppendorf tube and 25 µL detergent NP is added. The lysate was vortexed for 30 seconds at full speed, then centrifuged at 10000 g for 5 minutes. The supernatant was discarded, and the pellet resuspended in 4% PFA for 5 minutes before centrifugation at the same speed and time. The pellet was then resuspended in PBS and put through the 10 µm pluriStrainer cell strainer (PluriSelect; #43-50005-03). The flow-through was filtered through the 5 µm, pluriStrainer cell strainer (PluriSelect; #43-50005-13) and the final eluate was collected for sorting via the CellenONE microfluidics device.

The eluate was mapped on the CellenONE (CellenIon) to determine gating and particle size, with further dilutions performed as needed. Objects between 4 – 8 µm were considered to be MN and were sorted either singly or in groups of 10 (termed minibulks) for sequencing. Both DAPI and cGAS-GFP fluorescence were measured on the CellenONE but were not used as sorting parameters.

For the serial dilution experiments, RPE1 WT cells were harvested after nocodazole washout and the micronuclei purified as described previously. Instead of using the CellenONE, the final eluate was diluted as follows: 1/10, 1/100, and 1/1000 in sterile PBS. 1 μL of each concentration, including the undiluted eluate, was instead seeded directly into 96 wells of a 384-well sequencing plate with pre-dispensed primers. All lysis and library preparation steps were identical.

### Immunofluorescence

Cells were grown onto 22 x 22 mm coverslips in 6-well plates and fixed in PTEMF (0.2% Triton X-100, 0.02 M PIPES [pH 6.8], 0.01 M EGTA, 1 mM MgCl_2_, and 4% formaldehyde) for 10 minutes at room temperature. After fixation, coverslips were washed with PBS and then blocked in 3% BSA for 30 mins. Primary antibody incubation was carried out for 2 hours in blocking buffer with the following: anti-centromere (Antibodies Inc.; #15-234), and anti-phospho-Histone H2A.X (Sigma-Aldrich; #05-636). Coverslips were then washed three times with PBS and incubated with DAPI (Roche) and secondary antibodies in blocking buffer: goat anti-human AF647 (Stratech; # 109–606-088-JIR) and goat anti-mouse AF594 (Invitrogen; #A11005). Cover slips were then mounted on slides with Vectashield (Vector Laboratories; #H-1900-10), sealed with nail varnish, and imaged. All images were acquired on an Olympus DeltaVision RT microscope (Applied Precision, LLC) equipped with a Coolsnap HQ camera. Three-dimensional image stacks were acquired in 0.2 μm steps, using Olympus x100, x60, or x20 UPlanSApo oil immersion objectives. Image analysis was performed primarily using Softworx Explorer for micronucleus quantification and characterisation.

### Fluorescence *In-Situ* Hybridisation

Cells were grown directly onto imaging slides in 4-chambered dishes until 90% confluent. They were then fixed in 3:1 methanol:acetic acid for 10 minutes at room temperature. The slides were dehydrated in a graded ethanol series of 70%, 90%, and 100% for two minutes each, and stained with chromosome-specific centromere probes (CytoCell; LPE 001G, LPE 002R, LPE 007R). Slides and probe were denatured for 2 minutes at 75°C, then left overnight in a humid chamber at 37°C. The next day, they were washed in 0.25x SSC (Sigma-Aldrich; #S6639) for 2 minutes at 72°C, drained, and washed once more in 2x SSC and 0.05% Tween-20 (Sigma-Aldrich; #P1379) at RT for 30 seconds before DAPI staining and mounting.

### Sequencing and bioinformatics analysis

Single or minibulked MN were dispensed into 384 Lo-bind DNA plates containing unique pairs of Nextera i5 and i7 primers using the CellenOne microfluidics platform. Library preparation followed the tagmentase-based DLP+ method previously described (Zahn et al., 2019). Bioinformatics analysis was performed on the Queen Mary University of London (QMUL) High Performance Computing Cluster. Demultiplexed reads were trimmed using Trimmomatic using Nextera adaptors (Illumina) and length quality parameters and quality control was performed using FastQC. Trimmed reads were aligned to the GRCh38 genomic reference using BWA-MEM, with only reads over a map quality of 10 used for downstream analysis. The QDNAseq R package was used to bin aligned reads using 500 kb bins followed by GC correction and blacklisting. Log2-transformed read count values were extracted, and a threshold of 0.02% of total reads was applied to distinguish genuine signal from background noise. ACE was used to extract copy number segments. The data was further manually curated for quality control prior to scoring frequencies. All DNA fragments smaller than 5 Mb were excluded from analysis due to limitations of our sequencing resolution.

### Preparation of data and figures

All graphs were prepared and statistical testing conducted in the Prism software (version 11.0, GraphPad). All figures were assembled in Affinity Studio.

## Acknowledgements

We would like to dedicate this manuscript to the memory of Dr Siegfried Hauch (Qiagen). We thank the inspiring enthusiasm and highly enjoyable scientific discussions that were had throughout the early stages of this project with Siggi, and he is much missed. We would also like to thank the Henssen and Papathanasiou labs for sharing unpublished data and enjoyable scientific discussions. We would also like to thank members of the McClelland lab for critical reading of the manuscript.

