## Supplementary Figure 1 for "Rapid Micronucleus-Sequencing (RaMN-Seq) Reveals Patterns of Bias in Induced and Spontaneous Chromosomal Instability"

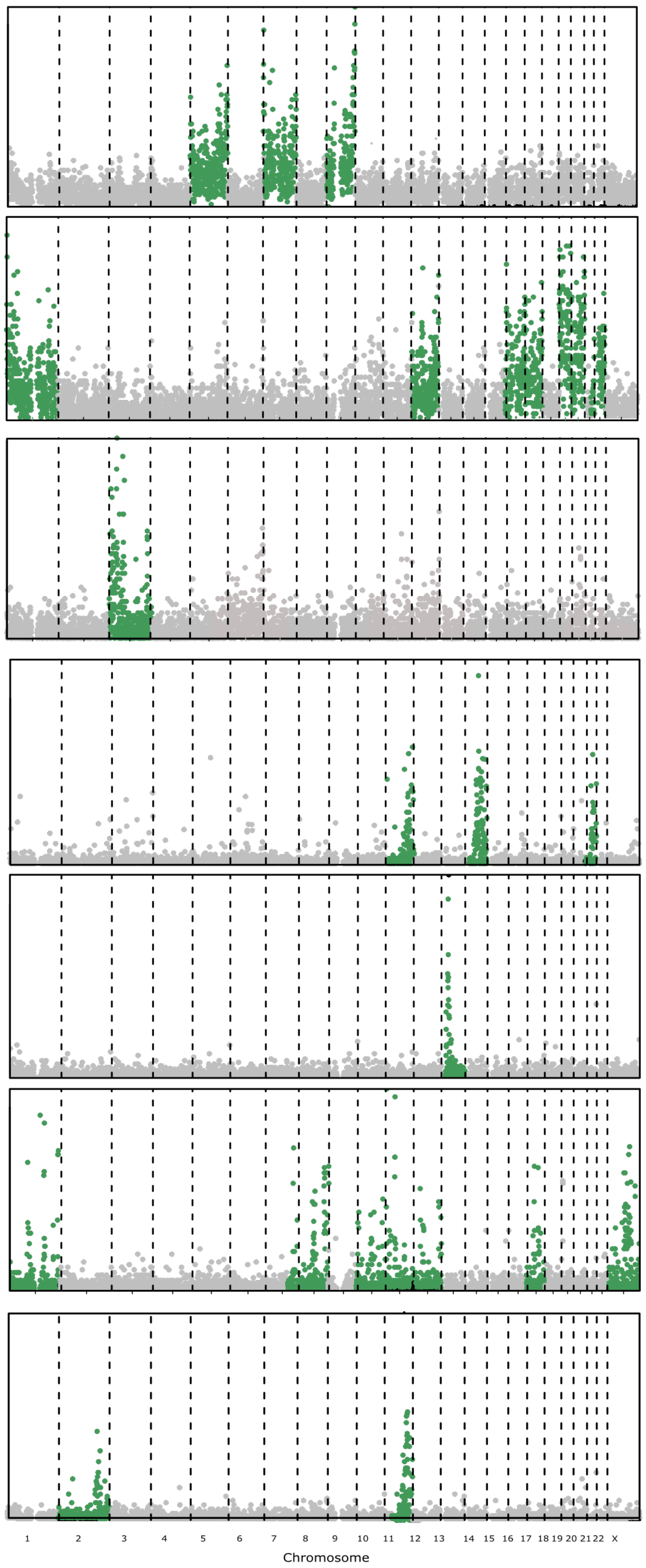


**Supplementary Figure 1: Dilution seeding allows single microcnucleus sequecning.** RPE1 wt cells were treated with nocodazole washout and subjected to RaMN-Seq. At the final eluate stage, serial dilutions were made across a 384 well plate as follows: undiluted; 1/10, 1/100, 1/1000. Micronuclei were detected at all dilutions but were most common in 1/10 and 1/100 dilutions. Nuclei were observed in the undiluted fraction but very rare in the other dilutions. Representative micronuclei are shown from across all dilutions.
